# ChIPbinner: An R package for analyzing broad histone marks binned in uniform windows from ChIP-Seq or CUT&RUN/TAG data

**DOI:** 10.1101/2024.10.02.616288

**Authors:** R. Padilla, E. Bareke, B. Hu, J. Majewski

## Abstract

**Background:** The decreasing costs of sequencing, along with the growing understanding of epigenetic mechanisms driving diseases, have led to the increased application of chromatin immunoprecipitation (ChIP-Seq), Cleavage Under Targets & Release Using Nuclease (CUT&RUN) and Cleavage Under Targets and Tagmentation (CUT&TAG) sequencing – which are designed to map DNA or chromatin-binding proteins to their genome targets – in biomedical research. Existing software tools, namely peak-callers, are available for analyzing data from these technologies, although they often struggle with diffuse and broad signals, such as those associated with broad histone post-translational modifications (PTMs).

**Results:** To address this limitation, we present ChIPbinner, an open-source R package tailored for reference-agnostic analysis of broad PTMs. Instead of relying on pre-identified enriched regions from peak-callers, ChIPbinner divides (bins) the genome into uniform windows. Thus, users are provided with an unbiased method to explore genome-wide differences between two samples using scatterplots, principal component analysis (PCA), and correlation plots. It also facilitates the identification and characterization of differential clusters of bins, allowing users to focus on specific genomic regions significantly affected by treatments or mutations. We demonstrated the effectiveness of this tool through a case study assessing H3K36me2 depletion following NSD1 knockout in head and neck squamous cell carcinoma, highlighting the advantages of ChIPbinner in detecting broad histone mark changes over existing software.

**Conclusions:** Binned analysis provides a more holistic view of the genomic landscape, allowing researchers to uncover broader patterns and correlations that may be missed when solely focusing on individual peaks. ChIPbinner offers researchers a convenient tool to perform binned analysis. It improves on previously published software by providing a clustering approach that is independent of each bin’s differential enrichment status, while also offering additional features for downstream analysis of these differentially enriched bins.

## Background

Histone modifications refer to chemical modifications to the histone tails of nucleosomes, the protein complexes around which DNA is wrapped. These modifications, also known as histone marks, regulate gene expression by facilitating or hindering access to the DNA. Narrow histone marks, such as acetylation of lysine 27 at histone 3 (H3K27ac) are deposited in specific, focused genomic regions whereas broad histone marks, such as methylation of lysine 36 at histone 3 (H3K36me), are diffused across the genome, covering large genomic domains. Recent advancements and decreasing costs of sequencing technology, such as ChIP, CUT&RUN and CUT&TAG sequencing, which allow mapping of protein-DNA interactions, for example histone marks, have led to their increasing application in biomedical research. The widespread adoption of these technologies is further driven by our growing understanding of epigenetic mechanisms driving disease and cancer.

Model-based Analysis of ChIP-Seq (MACS) is a widely used tool for identifying genomic regions where the target ChIP-Seq signal is enriched relative to background noise from a control DNA input or non-targeting antibody experiment [1]. Commonly known as a “peak-caller”, MACS was originally designed to detect transcriptional factor binding sites, which are found in highly specific genomic regions [1]. Detection of diffuse enrichment covering extended genomic regions often suffers from high noise level and lack of saturation in sequencing coverage [2]. Thus, peak-callers designed for broad histone marks have been developed, such as EPIC2 for ChIP-Seq data [3], the “--broad” feature in MACS, and Sparse Enrichment Analysis for CUT&RUN (SEACR) [4].

However, despite these advances in peak calling strategies for broad histone marks, there is often a discordance amongst peak callers as to what constitutes true signal enrichment. Furthermore, diffuse, broad domains become fragmented into smaller, often biologically meaningless peaks. This situation is further confounded when the nature of the genomic distribution of chromatin modifications changes: H3K27me3 in the presence of the pediatric glioma H3K27M (histone 3 lysine-to-methionine mutation at position 27) mutation from a broad to promoter-focused distribution [5] or H3K36me2 deposited by NSD1/2 (broad), versus NSD3/ASH1L (narrow), making it difficult to apply a uniform approach for comparative analysis [6].

As an alternative to being limited to some set of pre-defined regions (peaks), the genome can be divided into uniform windows, which can be referred to as “binning the genome”. The advantage of this strategy compared to peak-calling is that it is reference-agnostic – the analysis is unbiased without a prior set of references needed. Bins that are differentially enriched would be identified, but it does not apply prior assumptions or biases to those bins, as opposed to peak callers which rely on an algorithm and prior assumptions. Here, we present ChIPbinner, an R package designed to compare genome-wide changes in broad histone marks between two samples without relying on pre-identified regions. It can be used to normalize raw signal that has been binned in uniform windows and combine replicates. Furthermore, exploratory analyses can be conducted: the distribution of bins in genic and intergenic regions can be assessed, and genome-wide profiles of samples can be evaluated using PCA or correlation plots to assess consistency of replicates and separation of samples according to treatment. A primary motivation for performing ChIP/CUT&RUN/TAG sequencing is to identify genomic regions where significant changes occur between different treatment conditions, also known as differential binding (DB) sites. This can uncover potential mechanisms through which changes in binding may contribute to the treatment effect.

Thus, in addition to identifying clusters of bins that remain unchanged between different treatment conditions, ChIPbinner can also identify clusters of bins that exhibit differential changes and assess whether these clusters are enriched or depleted in specific classes of functionally annotated regions.

### Implementation

ChIPbinner is an open-source R package that can be installed using ‘remotes::install_github(“padilr1/ChIPbinner”)’ (latest version) in R. ChIPbinner has been written to minimize the number of external dependencies required for installation.

### Data pre-processing and input

ChIPbinner takes ChIP-Seq or CUT&RUN/TAG data binned in uniform windows in a BED format as input. Hence, users would need to firstly convert their aligned sequence reads in BAM format into BED format. One of the published tools that than can be used to perform this task is “bedtools *bamtobed*” [7].

### Normalization and transformation into bigWig

Next, users can normalize the raw read counts in their BED file to adjust for library size and, in the case of ChIP-Seq data, background signal. Furthermore, to allow for direct quantitative comparisons and to correct for other confounding experimental variables, such as variations in genome fragmentation and immunoprecipitation efficiency, users can quantitatively scale the histone mark signal using techniques such as ChIP with reference exogenous genome (ChIP-Rx) [8] or the genome-wide modification percentage information obtained from mass spectrometry, as described in Farhangdoost *et al*. (2021) [9]. Additionally, users have options to exclude artifact signals from the ENCODE blacklist regions [10] and filter out bins with low raw read count. The normalized and scaled signals are outputted in a bigWig track format, which is useful for dense, continuous data.

### Handling replicates and global sample profiles

To ensure replicates for a given condition cluster together, users can generate plots of PCA and/or correlation matrix-based hierarchical clustering using the normalized bigWig files from the previous step. These plots also offer insights as to the effect of a treatment or mutation on the global profiles of samples for a given histone mark. If satisfied with the consistency of replicates, the users can proceed to merge replicates, which takes the average signal per bin between replicates.

### Genic/intergenic scatterplot

Comparing two bigWig files, typically the treated versus untreated samples, users can generate a scatterplot for bins annotated as genic or intergenic. Genic regions were taken as the union of any intervals having the “gene” annotations in the curated Ensembl database [11] and intergenic regions were thus defined as the complement of genic ones. Annotated genic and intergenic regions are available for the following assemblies: mm10 (mouse) and hg38 (human).

### Identifying and annotating clusters

ChIPbinner utilizes HDBSCAN, a density-based hierarchical clustering method, to identify clusters of similarly-behaving bins [12]. HDBSCAN builds a comprehensive density-based clustering hierarchy, from which a simplified hierarchy containing only the most significant clusters can be easily extracted [12]. Users can specify the stringency of the clustering algorithm, such as the minimum size grouping for a cluster and how conservative the clustering will be – the more conservative, the more points will be declared as noise and clusters will be restricted to progressively more dense areas. Each cluster will be assigned a letter (A to Z) for reference in downstream analysis.

### Density-based plots

Density-based scatterplots offer a genome-wide overview of clusters that show enrichment, depletion, or remain unchanged between two samples for a specific histone mark.

### Enrichment and depletion analysis

After identifying and annotating clusters of bins, a Fisher’s exact test can be used to determine whether a specified cluster of bins significantly overlap a specific class of annotated regions against a background of all bins, as implemented in LOLA [13].

Additionally, the user can evaluate exclusively a specified cluster of genic or intergenic bins overlapping a specific class of annotated regions. In these cases, the background is stratified to only genic or intergenic regions to avoid spurious associations to annotations confounded by their predominantly genic or intergenic localization. Three curated datasets of annotated regions have been provided with the ChIPbinner package: Ensembl regulatory build [11], repeatMasker (www.repeatmasker.org) and ENCODE candidate cis-regulatory elements [14], which can be found at https://github.com/padilr1/ChIPbinner_database. Optionally, the user can provide their own dataset of annotated regions to test their cluster of bins against.

## Results and discussion

### Comparison with other software that identifies DB sites

Software tools for detecting DB sites in ChIP-Seq data have been developed using DiffBind [15] and csaw [16]. However, DiffBind relies on peak-sets derived from peak-callers to identify DB sites between sample groups [15]. Consequently, there is a lack of independence between peak calling and DB detection – DiffBind is constrained by the same assumptions and biases associated with peak-callers. Conversely, csaw uses a window-based strategy to summarize read counts across the genome and is independent of peak-callers. It uses statistical methods in the edgeR package, which was designed for differential gene expression analysis [17], to test for significant differences in each window. Finally, it clusters windows into regions and controls the false discovery rate over all detected regions [16]. However, the default clustering procedures for csaw relies on independent filtering to remove irrelevant windows. This ensures that the regions that are differentially changing are reasonably narrow and can be easily interpreted, which is typically the case for TFs and narrow histone marks.

However, enriched regions tend to be very large for more diffuse marks and such regions may be difficult to interpret when smaller DB subintervals is of interest and not necessarily well-separated, defined regions. To address this issue, csaw employs a post-hoc analysis for diffuse histone marks, using only significant windows clustering [16]. This approach defines and clusters significant windows while controlling the cluster-level FDR at 5%. This alleviates the need for stringent abundance filtering to achieve well-separated regions prior to clustering for diffuse marks. However, the calculation of the cluster-level FDR is not entirely rigorous, and the clusters are influenced by the DB status of the windows. Thus, csaw struggles to detect DB properly when dealing with the diffuse signal from broad histone marks. Overall, although csaw avoids the aggressive assumptions when defining peaks as no peak calling is performed, it introduces other assumptions by applying a statistical algorithm to detect differentially bound sites prior to clustering. Furthermore, important, but subtle biological differences might be missed when the statistical power is too low, which often occurs with a limited number or lack of replicates.

In contrast to csaw, ChIPbinner does not rely on a statistical algorithm. The clustering of bins is independent of the DB status of each bin - normalized read counts per window from the two samples are directly inputted into the clustering algorithm without any prior statistical comparisons. Furthermore, although designing sequencing experiments without replicates is discouraged, ChIPbinner can be used with only a single replicate per treatment, allowing cross-validation across cell lines as independent controls. ChIPbinner includes additional features that are not readily available in csaw, such as built-in functions for PCA, hierarchical clustering, annotating individual bins as genic or intergenic, and enrichment/depletion analysis for specified clusters. Lastly, ChIPbinner offers functionalities that enable users to easily normalize and quantitatively scale the signal in their bins, whereas csaw requires some manual coding to achieve the same results.

### Case study

The methods implemented in ChIPbinner are largely derived from our analyses in Farhangdoost et al. (2021), where we identified and characterized genomic compartments that exhibited the greatest loss of the broad histone mark H3K36me2 following NSD1 knockout (NSD1-KO) in a head and neck squamous cell carcinoma (HNSCC) cell line Cal27. In that study, we subdivided the genome into 10-kilobyte bins. Using an approach similar to what can be performed with ChIPbinner, we identified a cluster of bins, designated as Cluster B, with high initial levels of H3K36me2 in the parental cell lines (WT) and low levels in the NSD1-KO (Fig. 2a). For comparison, we used csaw with 10kb bins and cluster-level FDR at 5% to detect a similar cluster of bins with downregulated levels of H3K36me2 in NSD1-KO compared to WT (Fig. 2b, 2c; purple boxes). 86 % (51607/59848) of downregulated DB bins identified using csaw were also detected by ChIPbinner whereas csaw only detected 55 % (51607/93613) of the DB bins identified by ChIPbinner (Fig. 2c). Thus, compared to csaw, ChIPbinner seems to perform better in capturing differential changes in the broad histone mark H3K36me2 following NSD1-KO in HNSCC.

**Fig. 1.**
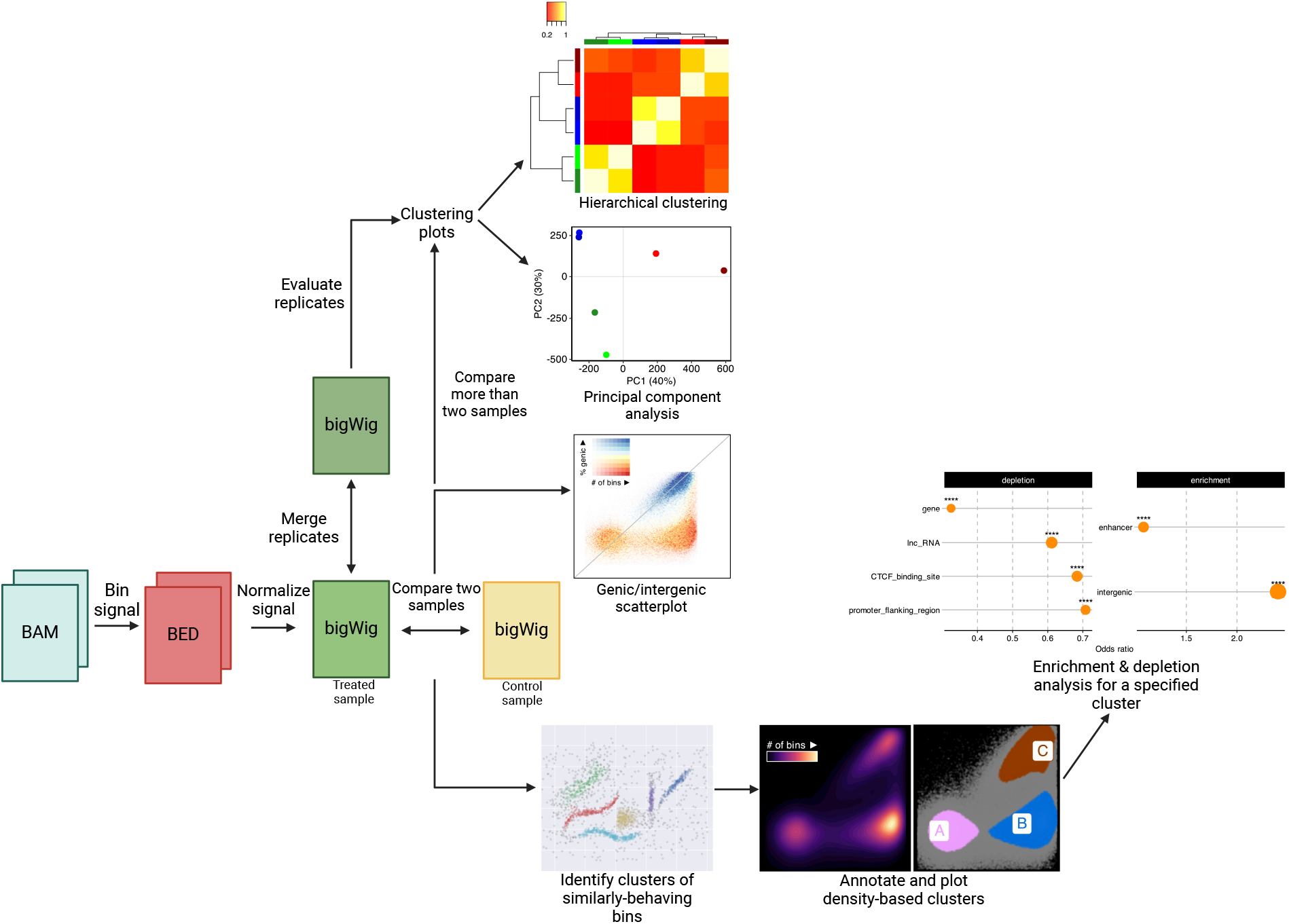
The ChIPbinner workflow requires that aligned reads from ChIP-Seq or CUT&RUN/TAG experiments, typically in BAM format, to be binned in uniform windows and converted to BED (Browser Extensible Data) format (minimum BED3 format with chromosome, start and end information) (red box). Users can normalize the raw binned signal. Pairwise comparisons, such as treated versus control (yellow box), can then be performed using genic/intergenic scatterplots, which annotates each bin as genic or intergenic. For comparing more than two samples, PCA or hierarchical clustering can be used to assess consistency of replicates or effect of treatment/mutation on genome-wide profiles of each sample. Next, users can combine replicates for use in downstream analysis (green boxes). After selecting two samples for pairwise comparison, users can identify and annotate clusters of similarly-behaving bins and visualize them. Finally, specific clusters of bins can be selected for enrichment and depletion analysis.

**Fig. 2.**
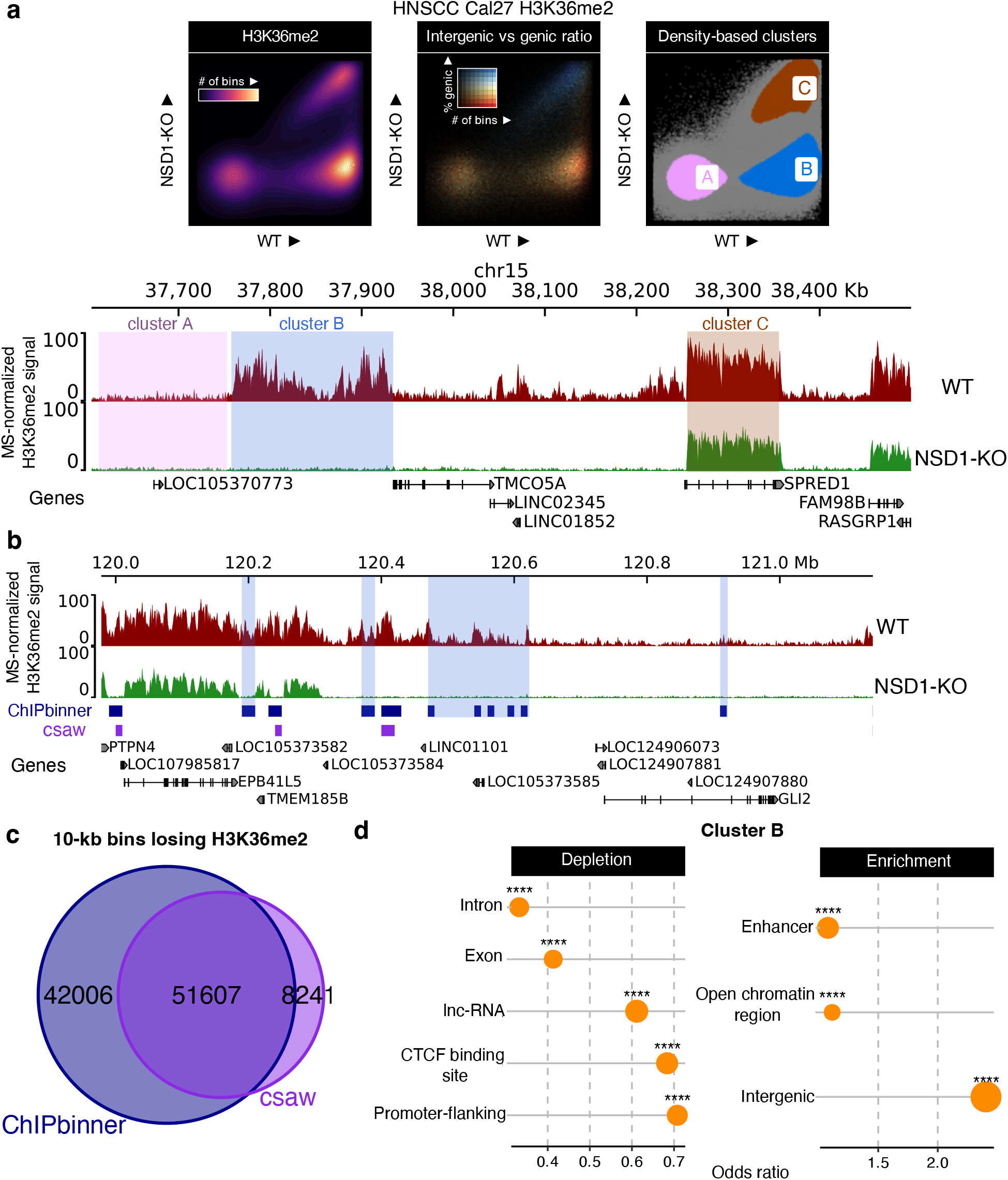
A case study in a head and neck squamous cell carcinoma (HNSCC) cell line Cal27, which belong to the HPV(-) subgroup of HNSCC but harbor no endogenous mutations affecting H3K36me. **a**. Scatterplots and genome-browser tracks demonstrating how clusters of 10-kilobyte (kb) bins identified using ChIPbinner correspond to specific genomic regions. A comparison of the wildtype cell line (WT) (dark red) and an NSD1 knockout cell line (NSD1-KO) (green) reveals depletion of intergenic H3K36me2 in Cluster B (highlighted in blue), where high levels of H3K36me2 are present in WT and low levels are found in NSD1-KO. In clusters A (highlighted in pink) and C (highlighted in brown), there is low and high levels of H3K36me2 in both conditions respectively. ChIP-seq signals were normalized using genome-wide modification percentage values obtained from mass spectrometry (MS). **b**. Genome-browser tracks showing an example locus where ChIPbinner (blue boxes) detected more regions of H3K36me2 loss compared to csaw (purple boxes). **c**. Genome-wide overlap analysis revealed that out of the 10-kb bins with loss of H3K36me2, ChIPbinner detected 51,607 out of 59,848 (86%) bins identified by csaw, while csaw only detected 51,607 out of 93,613 (55%) bins identified by ChIPbinner. **d**. ChIPbinner allows users to perform differential enrichment and depletion analysis for a specified cluster. In this case, cluster B bins were tested for overlap with a specific class of annotated regions from Ensembl. **** represents p-value < 0.0001 based on Fisher’s exact test. The results show that cluster B bins are enriched in intergenic regions, particularly at enhancers and open chromatin regions, and depleted in promoter flanking regions, exons and introns.

Nevertheless, as stated previously, ChIPbinner also allows users to perform downstream analysis following identification of clusters. In our case study, characterization of Cluster B indicated that these regions were predominantly intergenic and enriched for enhancers (Fig. 2a,d) [9] – both of which can be determined for a specific cluster using the genic/intergenic scatterplot and enrichment/depletion functions, respectively, available in the ChIPbinner package. Further characterization of cluster B revealed that these regions exhibited the greatest reduction in H3K27ac binding, and led to the strongest downregulation of target genes within the same topologically associating domain (TAD) [9]. More recently, we used ChIPbinner to identify genomic regions with significant loss of the broad, heterochromatic histone mark H3K9me3 following complete depletion of H3K36me in mouse mesenchymal stem cells [18]. These regions were predominantly intergenic, and we observed significant upregulation of genes and transposable elements within them. We replicated this binned analysis in HNSCC cell lines, where we similarly identified a cluster of bins significantly losing H3K9me3 following H3K36me depletion [18]. In another study from our group, we used intergenic/genic scatterplots for binned genomic regions—as can be generated with ChIPbinner—to demonstrate that the greatest reduction in H3K36me2 following NSD1 knockout in mouse embryonic stem cells occurs primarily at intergenic regions [19]. These studies highlight the value of ChIPbinner in analyzing broad histone marks, such as H3K36me2 and H3K9me3, for identifying and characterizing genomic compartments undergoing the most significant changes, or as a visualization tool for genome-wide comparisons between different samples.

## Conclusions

ChIPbinner offers an alternative approach to peak-callers and equips researchers with an unbiased tool to analyze trends and patterns in epigenetic data, especially for broad histone marks. By providing a compiled workflow and dedicated functions, our software saves researchers time and effort in implementing binned analysis techniques. It also ensures that their analyses can be easily replicated and applied to new datasets, even for users with limited programming experience. ChIPbinner improves on previously published software by offering a clustering approach that is independent of DB and including additional features for downstream analysis.

In systems characterized by large global changes in the distribution or levels of broad histone marks, as observed here following NSD1-KO for H3K36me2 and NSD1/2-SETD2-TKO for H3K9me3, ChIPbinner is a useful tool for exploring differences of chromatin modifications between conditions. It allows users to analyze changes in the distribution of broad histone marks within genic and intergenic regions, without the assumptions or constraints imposed by peak callers or stringent statistical algorithms.

### Availability and requirements

**Project name:** ChIPbinner

**Project home page:** https://github.com/padilr1/ChIPbinner

**Operating system:** Platform independent

**Programming language:** R

**Other requirements:** None

**License:** GPL-3.0

**Any restrictions to use by non-academics**: None

## Abbreviations

ChIP-Seq: Chromatin immunoprecipitation sequencing
CUT&RUN: Cleavage Under Targets & Release Using Nuclease
CUT&TAG: Cleavage Under Targets and Tagmentation
PTM: Post-translational modifications
PCA: Principal component analysis
H3K27ac: Histone three lysine 27 acetylation
H3K36me: Histone three lysine 36 methylation
MACS: Model-based Analysis of ChIP-Seq
SEACR: Sparse enrichment analysis for CUT&RUN
H3K27M: Lysine-to-methionine mutations in histone 3 at position 27
NSD: Nuclear receptor-binding SET domain protein
ASH1L: Absent, small, or homeotic discs 1-like
DB: Differential binding
BED: Browser extensible data
ChIP-Rx: ChIP with reference exogenous genome
KB: Kilobyte
WT: Wildtype
NSD1-KO: NSD1 knockout
HNSCC: head and neck squamous cell carcinoma
TAD: Topologically associating domain

## Declarations

### Ethics approval and consent to participate

Not applicable.

### Consent for publication

Not applicable.

## Availability of data and materials

The datasets analysed during the current study are available in the National Center for Biotechnology Information Gene Expression Omnibus (NCBI-GEO) under accession number GSE149670, https://www.ncbi.nlm.nih.gov/geo/query/acc.cgi?acc=GSE149670 [9]. Curated databases from Ensembl, RepeatMasker and ENCODE candidate cis-regulatory elements can be found at https://github.com/padilr1/ChIPbinner_database.

## Authors’ contributions

R.P. conceptualized the software tool with the help of J.M. B.H. provided the initial scripts. R.P. developed and compiled software package, with assistance from E.B. R.P. wrote the initial manuscript, with suggestions and contributions from J.M., E.B. and B.H. for the final manuscript. All authors read and approved the final manuscript.

